# The ‘breakpoint’ in transmission for soil-transmitted helminths: dynamical behaviour near the equilibria and the impact of infected human migration

**DOI:** 10.1101/619908

**Authors:** Robert J. Hardwick, Carolin Vegvari, James E. Truscott, Roy M. Anderson

## Abstract

Building on past research, we here develop an analytic framework for describing the dynamics of the transmission of soil-transmitted helminth (STH) parasitic infections near the transmission breakpoint and equilibria of endemic infection and disease extinction, while allowing for perturbations in the infectious reservoir of the parasite within a defined location. This perturbation provides a model for the effect of infected human movement between villages with differing degrees of parasite control induced by mass drug administration (MDA). Analysing the dynamical behaviour around the unstable equilibrium, known as the transmission ‘breakpoint’, we illustrate how slowly-varying the dynamics are and develop an understanding of how discrete ‘pulses’ in the release of transmission stages (eggs or larvae, depending on the species of STH), due to infected human migration between villages, can lead to perturbations in the deterministic transmission dynamics. Such perturbations are found to have the potential to undermine targets for parasite elimination as a result of MDA and/or improvements in water and sanitation provision. We extend our analysis by developing a simple stochastic model and analytically investigate the uncertainty this induces in the dynamics. Where appropriate, all analytical results are supported by numerical analyses.

## 1. Introduction

The past two decades have seen considerable progress in the control of many of the Neglected Tropical Diseases (NTDs) in resource-limited countries [1, 2, 3]. This is especially the case for helminth parasites where drugs are available to treated infection via mass drug administration (MDA) programmes.
These infections include the soil-transmitted helminths (STH), the filarial worms and the schistosome parasites. Post the London Declaration in 2010, pharmaceutical companies have donated drugs free to resource poor countries with endemic infection to facilitate large mass drug administration (MDA) control programmes under the direction of World Health Organization (WHO) guidelines for treatment strategies in various transmission settings [2, 4, 5].

These important helminth parasites — that are a cause of much morbidity and, in some cases, mortality — do not induce strong acquired immunity to reinfection post anthelmintic treatment. MDA must therefore be repeated frequently to reduce morbidity where the frequency of treatment depends on the intensity of transmission in a defined setting (the magnitude of the basic reproductive number *R*_0_) and other factors associated with the population biology of the parasite such as adult parasite life expectancy [6]. In these circumstances, recent research has focused on the question of whether MDA (if administered at high coverage, frequently and targeted at large sections of the community) can, on its own, eliminate helminth parasite transmission [7, 8, 9, 10]. For STH, a number of large-scale trials are currently underway in regions of endemic infection to test this notion [11, 12].

Mathematical models of helminth parasite transmission and MDA effect provide many insights into both the overall impact of various control policies and the behaviour of parasite populations under sustained drug treatment at defined levels of population coverage and treatment frequencies [9, 10]. Helminths are dioecious by nature having separate sexes. As such, both male and female parasites must be in the same host for the female worm to be fertilized and produce viable infective stages (eggs or larvae) to sustain transmission. Past research has shown that the parasite population has three possible equilibria mean worm loads in the human host, a stable endemic state, parasite extinction, and an unstable state termed the ‘transmission break-point’ which lies between the stable state and parasite extinction [13, 14, 15]. The distance between the unstable breakpoint and parasite extinction is determined by the degree of parasite aggregation accross individuals, as measured inversely by the negative binomial shape parameter *k*, where high aggregation draws the break-point towards the point of parasite extinction [6].

The past successes of MDA control programmes have now moved parasite mean intensities and prevalences of infection to low levels in many regions of endemic infection in Africa and Asia. This has provided a further stimulus to push to eliminate parasite transmission where possible by increased efforts to raise coverage and compliance to treatment to remove the need for continued MDA programmes into the predictable future. In these circumstances, a deeper understanding of a number of epidemiological factors is ideally required and these include the two issues addressed in this paper.

The first of these factors is the expected dynamical behaviour of the human host-helminth parasite system around the break-point in transmission, which separates between the two dynamical attractors of endemic parasite persistence and parasite extinction. An obvious question is: how slowly (or quickly) do the parasite transmission dynamics move away from the unstable breakpoint towards either attractor? This question is of practical significance since in monitoring and evaluating parasite prevalence and mean intensity trends as MDA coverage increase, public health workers need to understand how these epidemiological measures might be expected to change over time as the system moves towards parasite transmission interruption.

The second factor is the influence of the migration of infected individuals into areas where the parasite population is set to cross the boundary of transmission cessation. This issue arises due to the observed heterogeneity in MDA coverage between adjacent or nearby villages or towns, due to various factors influencing the delivery and acceptance of MDA. Such migrations can potentially modify the reservoir of infectious material (eggs and larvae for STH) and hence render the transmission dynamics uncertain. This, in turn, poses the obvious question: how uncertain are the trajectories of transmission dynamics in the presence of migration? This issue is again of practical significance since an understanding of the impact of human movement between population centres in a landscape of heterogeneous MDA coverage will inform policy formulation and focus attention on attaining high and uniform coverage.

We examine both of these questions using mathematical models of parasite transmission and control and employing analytical and numerical approaches. Our focus is on the control of the soil-transmitted helminths, but the conclusions are more broadly applicable to other human helminth infections. Throughout our focus shall be on the applied significance of the predicted dynamical properties of human-helminth parasite interactions to the design of effective control policies and their monitoring and evaluation.

## 2. Basic transmission model

The nonlinear dynamical system of equations describing the time evolution of the mean total worm burden *M*(*t*) (hereafter, simply ‘mean worm burden’) and infectious reservoir *L*(*t*) is given by [6]

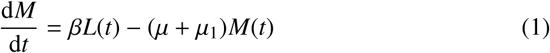

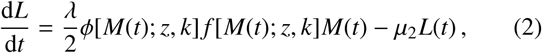

where: *β* quantifies the contact rate of an individual with respect to *L*; *µ, µ*_1_ and *µ* _2_ are the death rates associated to humans, worms and infectious material in the reservoir (eggs and larvae), respectively; *λ* is the rate of egg production per female worm; *γ* is the density-dependent fecundity power index which accounts for a decreased egg rate per female worm in the hosts with a large number of worms aggregated together; and we have defined *z* ≡ *e*^-*γ*^ as well as

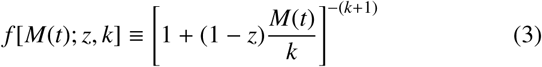

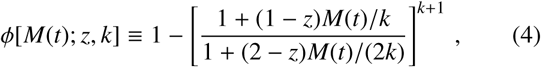

where the former factor quantifies the effect of density-dependent fecundity, the latter factor quantifies the effect of sexual reproduction between worms in order to generate new fertilized eggs for the reservoir (assuming fully polygamous male worms) and 1/*k* is the ‘clumping factor’ (the degree to which worms aggregate within individual hosts).

The second derivative of *M*(*t*) can be obtained by taking the time derivative of Eq. (1)

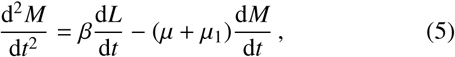

which is essentially a ‘force’ acting in the 4-dimensional phase space of [*M*(*t*), *L*(*t*), *M*′(*t*), *L*′(*t*)]. With a specified reservoir behaviour *L*(*t*), Eq. (1) and Eq. (5) hence describe the dynamics of *M*(*t*) towards stable ‘equilibria’ — the latter term corresponding to regions of the phase space where the first time derivatives *M*′(*t*) and *L*′(*t*) vanish.

### 2.1 Reservoir in equilibrium

We simplify the analyses by considering the case where the infectious reservoir of infection in the human habitat (eggs in the case of *Ascaris lumbricoides* or *Trichuris trichuria* — or larvae in the case of the hookworms: *Necator americanus* and *Ancylostoma duodenale*) is at equilibrium. This assumption is justified by the relative long life span of the adult worm in the human host (1 to 2 years) versus the life expectancy of the eggs or larvae outside the human host which is in terms of weeks to a few months — see Ref. [6]).

Consider now the initial situation where the reservoir is at equilibrium, hence a trivial manipulation of Eqs. (1) and (2) yields

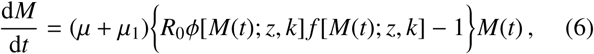

where, following the standard methodology, we have also collated many of the parameters into *R*_0_, the density effect-independent basic reproduction number^1^

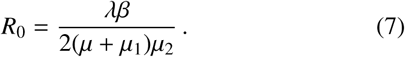

Note, by setting the reservoir at equilibrium, Eq. (5) implies also that

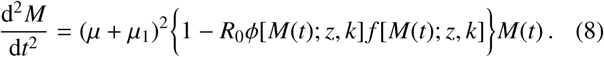

Given Eq. (6) we may thus read off the condition for equilibrium mean worm burden *M*_*_

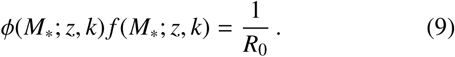

Eq. (9) may only be solved precisely by numerical approaches. Using a root-finding algorithm^2^ we plot an example of the equilibrium solutions given values of *k* = 0.3 and *γ* = 0.08 with the dashed and solid black lines in both panels of Fig. 1. For the purpose of consistency between plots, in Fig. 1 and throughout this work (unless otherwise explicitly stated) we shall make the following parameter choices: *µ* = 1/70 years^-1^, *µ*_1_ = 1/2 years^-1^, *µ*_2_ = 5 years^-1^, *λ* = 10, *β* = 1, *γ* = 0.08, *k* = 0.3 and, for this choice of parameters, Eq. (7) determines *R*_0_≃1.94.

**Figure 1.**
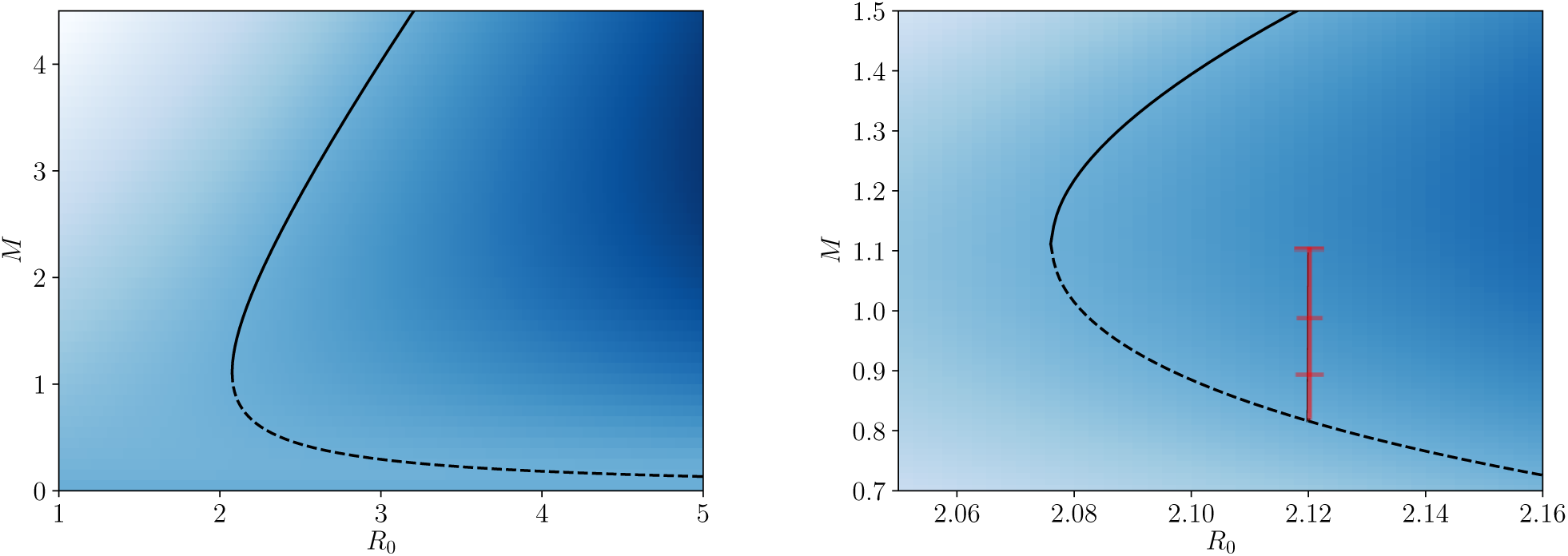
A visualisation of the phase plane generated by Eq. (6) (with *k* = 0.3 and *γ* = 0.08) where black solid and dashed lines correspond to the two branches of equilibrium solution numerically obtained by satisfying Eq. (9). In both panels the heatmap corresponds to the strength and direction of the first derivatives *M*′(*t*) (and second, up to a negative constant −(*µ* + *µ*_1_)*M*′(*t*) — see Eq. (8)) computed through Eq. (6) in the vertical direction (lines of constant *R*_0_), where lighter colours correspond to strongly negative values (strong forces downwards) and darker colours correspond to strongly positive values (strong forces upwards) — consequently, intermediate colours have the weakest forces in either direction. The right panel is a zoomed version of the left panel with enhanced colour contrast for illustration purposes. The increasing timescales to travel between distances of *M* = [1.0 → 1.1, 0.9 → 1.0, 0.83 → 0.9] for a fixed value of *R*_0_ = 2.12 are *t* −*t*_0_ ≃ [9.6 years, 12.9 years, 23.4 years], where we have indicated the trajectories by red markings.

Two equilibria are present in the solution to Eq. (9): the first is typically referred to as ‘stable’ (represented by the solid black line in Fig. 1) as this is the endemic solution to the STH epidemic which is an attractor for a range of values of *R*_0_ > *R*_0†_ (where *R*_0†_ refers to the value of *R*_0_ at which the two equilibria collide); and the second is typically referred to as ‘unstable’ (represented by the dashed black line in Fig. 1) as it corresponds to a repellor in the phase plane, i.e., a barrier where values of *M*(*t*) above it are attracted away to the stable equilibrium and values of *M*(*t*) below it are attracted away to the disease extinction equilibrium *M*(*t*) = 0 — the trivial solution to Eq. (6).

Two points are marked in the right panel Fig. 1 as an illustration of the lengths of time — or ‘timescales’ — involved for the motion of the mean worm load near the unstable breakpoint in transmission. Around the breakpoint the system may hover for tens of years (or even greater timescales) before either moving back to the stable state of endemic infection or the state of parasite transmission extinction (zero mean worm load). In practical terms this is an important observation. It suggests that as control measures intensify to move the system to the point of parasite transmission extinction, average worm loads may exhibit a long time (many years) approaching the state of parasite eradication.

The long time periods near the unstable breakpoint have important ramifications for ongoing monitoring and evaluation programmes: they imply that little progress may be observed in lowering parasite burdens before the control measures eventually ‘push’ the parasite population to extinction. The timescales of these movements will depend critically on the adult parasite life span in the human host, as well as the size of the *R*_0_ value. For STH, the right panel of Fig. 1 indicates in red the distances of *M* = [1.0 → 1.1, 0.9 → 1.0, 0.83 → 0.9], where for a fixed value of *R*_0_ = 2.12 one obtains timescales to travel these distances of *t*−*t*_0_≃ [9.6 years, 12.9 years, 23.4 years]. Timescales for STH hovering below the unstable breakpoint before extinction will take a similar value to the adult worm lifespan of 1-2 years, whereas, for Filarial worms with very long average lifespans in the human host of 7 years or more, this timescale may be decades.

### 2.2 Perturbing the system with migration

Notice that by perturbing the system away from reservoir equilibrium by δ*L*(*t*) = *L*(*t*) − *L*_eq_, and by approximating the dynamics of *M*(*t*) to be roughly constant over the time period that it takes *L*(*t*) to equilibriate,^3^ such that Eq. (2) becomes

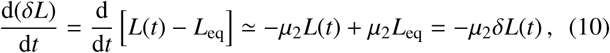

we are able to find a modification of Eq. (6) — by inserting the solution to Eq. (10) — which accounts for a perturbation in the infectious reservoir. Under such a perturbation, which may arise from the migration of people in or out of the spatial region represented by the infectious reservoir, Eqs. (6) and (8) become

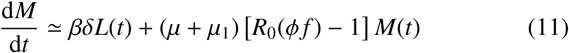

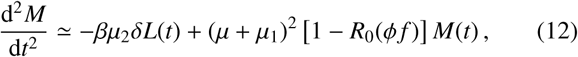

where we have defined the shorthand notation (ϕ *f)* ≡ϕ[*M*(*t*); *z, k*] *f* [*M*(*t*); *z, k*] and, with some initial time *t*_0_ set, we have used the approximate solution

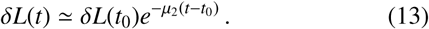

Eqs. (11) and (13) thus immediately indicate that a mechnism for fluctuation to or away from the equilibria in, e.g., Fig. 1, is possible and that a deeper study of the dynamics of the system is necessary in order to fully understand the ramifications.

## 3. Saddle-node bifurcation expansion

In this section, we shall introduce the concept of a ‘saddle-node bifurcation’, which is found to be present at the point when the two equilibria collide in, e.g., Fig. 1. This concept allows for the development of a formalism — which we broadly discuss here and in more derive with more detail in Appendix A — that provides a local approximative dynamical description of the human-helminth system.

We illustrate the point at which the stable endemic equilibrium (solid black line) and unstable breakpoint equilibrium (dashed black line) meet, known as a saddle-node bifurcation, in Fig. 1. In Appendix A we demonstrate how to obtain the value of the mean worm burden at this point *M* = *M*_†_(*z, k*), however here we shall simply quote it as

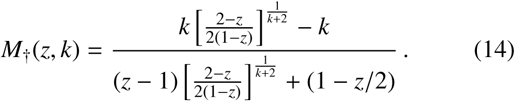

From this value, one may deduce the basic reproduction number at this point *R*_0_ = *R*_0†_(*z, k*), which we introduced in Sec. 1, by inverting Eq. (9) such that

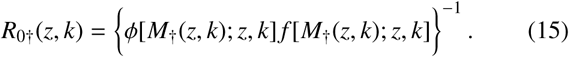

### 3.1 In the absence of migration

Given the existence of this saddle-node bifurcation, locally about *M*_†_(*z, k*) we may expand the system (suppressing dependencies and defining (ϕ *f)*_†_ ≡ϕ[*M*_†_(*z, k*); *z, k*] *f* [*M*_†_(*z, k*); *z, k*] for brevity) such that

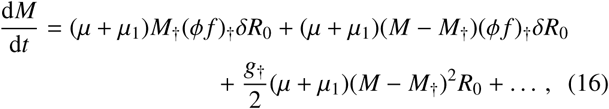

where we have defined δ *R*_0_ = δ*R*_0_(*z, k, R*_0_) ≡ *R*_0_ – *R*_0†_(*z, k*) and a new function

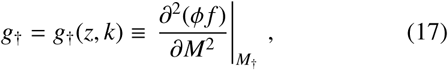

which is given explicitly in Appendix A. Note that, for typical values of *z*, the *g*_†_(*z k*) function takes negative values which decrease in magnitude with increasing *k*.

Now by truncating the expansion in Eq. (16), keeping up to 𝒪(*M*^2^) terms,^4^ we are able to solve the system exactly to find the following analytic solution

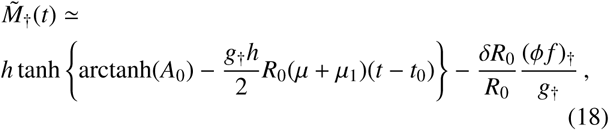

where 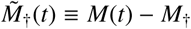 and we have defined

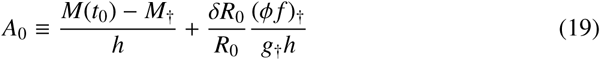

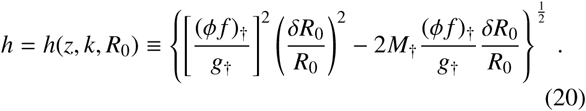

Note that when *k* ∼𝒪(0.1) or larger, our expansion in Eq. (16) is most accurate in the limit where |*M* −*M*_†_(*z, k*)| ≪ 1.^5^ In both panels of Fig. 2 we plot our approximative solution to the transmission dynamics given in Eq. (18) against the full solution to the dynamics obtained numerically,^6^ represented by the coloured solid and dashed lines, respectively.

**Figure 2.**
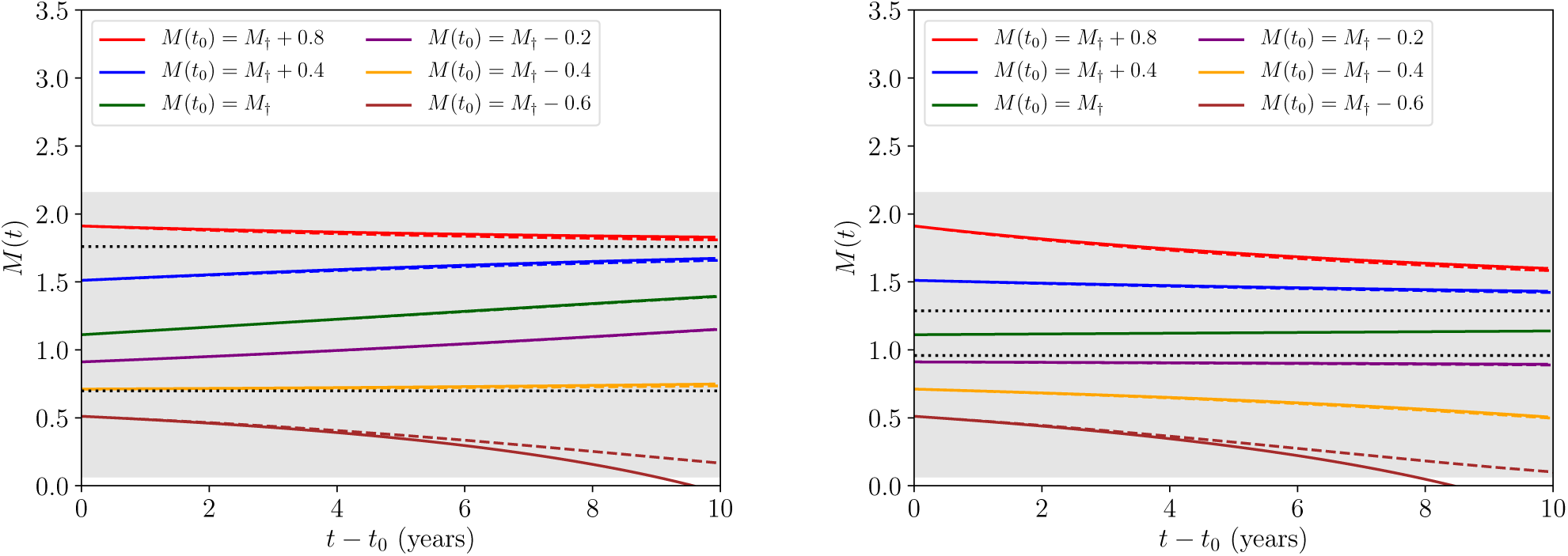
The approximative solution to the transmission dynamics given in Eq. (18) against the full solution to the dynamics obtained numerically, each represented by the coloured solid and dashed lines, respectively, for a range of initial conditions *M*(*t*_0_). In the left panel we have fixed a value of *R*_0_ = *R*_0†_(*z, k*)+ 0.1 ≃ 2.18 and in the right panel with a value of *R*_0_ = *R*_0†_(*z, k*)+ 0.01 ≃ 2.09. The dotted black horizontal lines corresponds to the value of *M* at the stable (upper) and unstable (lower) equilibria numerically obtained from Eq. (9). Lastly, the grey region corresponds to values for which |*M* -*M* _†_(*z, k*)|< 1 and hence the expansion used to obtain Eq. (18) leads to good agreement with the full numerical solution.

In the figure, it is also clear that the agreement between solutions is best *∀M*(*t*) (including the initial condition *M*(*t*_0_)) when the dynamics are confined within the grey region representing |*M* -*M* _†_(*z, k*)| < 1, supporting our expansion accuracy argumentation. The black dashed horizontal lines mark the value of *M* at the stable endemic equilibrium (upper) and unstable breakpoint equilibrium (lower) numerically obtained from Eq. (9), hence the full solutions, as well as those approximate solutions which initialise |*M* (*t*_0_)–*M*_†_(*z, k*)|< 1, are expected to change direction upon crossing the lower threshold.

Our analysis has now reached the point where we are in a position to address the initial question posed by the introduction. Moving from left to right between the two sets of panels in Fig. 2, one decreases in value of the basic reproduction number from *R*_0_ = *R*_0†_(*z, k*)+ 0.1 ≃ 2.18 to *R*_0_ = *R*_0†_(*z, k*) + 0.01 ≃ 2.09. Despite the effect this change has on the position of the equilibria, in both cases it is clear that as trajectories near the transmission breakpoint (the lower of the two dashed horizontal black lines) the rate of change in the transmission dynamics becomes extremely slow. This is most notable by considering that the timescale over which both panels are plotted is 10 years — a substantial period for the transmission dynamics to not vary significantly.

Another way to quantify the rate of change in the human-helminth system near the disease breakpoint is to compute the timescales over which the dynamics evolve. For completeness, we shall consider one such timescale directly in the next section.

### 3.2 Timescales away from the unstable equilibrium

In the previous sections we have been able to obtain approximate solutions to the dynamical behaviour parameterically near the point of saddle-node bifurcation. In contrast to this, if one wishes to compute the timescales towards fixed values over large regions of the phase plane, a numerical approach is necessary to avoid significant error.

In order to compute the time it takes for the transmission dynamics to travel between values of *M* = *M*_init_ and *M*_end_ in the absence of migration, for instance, one must compute the following integral

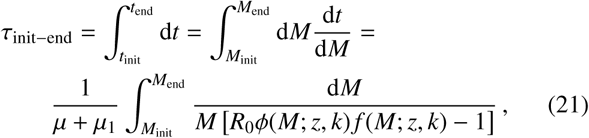

where Eq. (6) has been inserted into the expression in order to obtain the last equality. By way of example, in Fig. 3 we plot the numerically-obtained solutions to Eq. (21) for the length of time it takes to travel to *M* = *M*_†_(*z, k*) for different initial values and worm death rates *µ*_1_.

**Figure 3.**
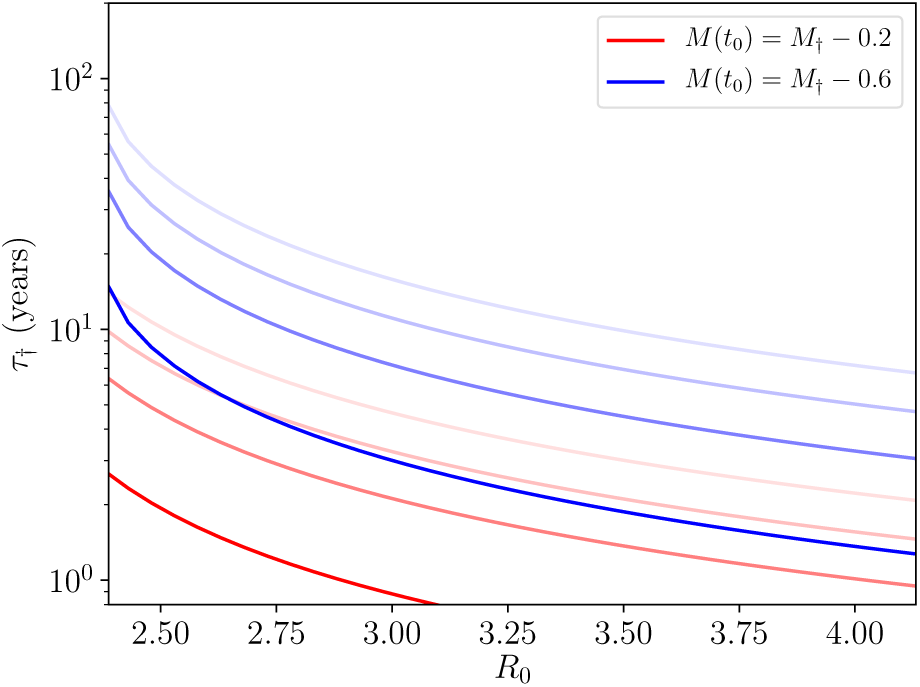
Numerically-obtained values using integral in Eq. (21) to compute the length of time it takes for the transmission dynamics to reach the value of *M* = *M* _†_(*z, k*) for two different initial values of *M* = *M*(*t*_0_), as shown in the legend. A range of worm death rates have been used of *µ*_1_ = 1/2 years^-1^, 1/5 years^-1^, 1/8 years^-1^, 1/12 years^-1^ in decreasing order, which corresponds to a fading colour in the plotted lines.

Fig. 3 is a further illustration of an essential point made at the end of the previous section: that the local transmission dynamics around the unstable breakpoint equilibrium are extremely slow so that the timescales away from this region lengthen as one nears it.

Motivated by the good agreement in the previous section between our analytic approximations and the full numerical solution to the STH transmission dynamics within a controlled range of initial conditions, we shall proceed to develop an equivalent approach with the inclusion of a perturbation from the infectious reservoir such that we may begin to address our second question.

### 3.3. Including a migration perturbation

Consider a perturbation of the form given by Eq. (13). We shall refer to this throughout this section as a ‘migration perturbation’ due to the equivalence between including the out-of-equilibrium dynamics of the infectious reservoir and the effective displacement provided by the variation in human population numbers. Given this perturbation, we have already obtained Eq. (11) to describe the transmission dynamics, however, as with Eq. (1), this equation also may only be solved precisely by numerical approaches in order to check the accuracy of our results.

By truncating an expansion of Eq. (11), keeping up to 𝒪(*M*^2^) terms in same way as in Eq. (16), we find that the resulting approximate equation takes a Riccati form

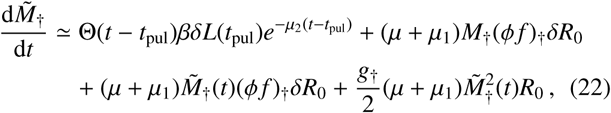

where, once again, 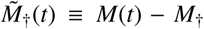 and Θ(*t* − *t*_pul_) is the Heaviside (step) function which initialises the migration pulse at time *t*_pul_ and *δL*(*t*_pul_) is the maximal amplitude of the perturbation in the infectious reservoir.

Remarkably, evolving the elapsed time from *t*_pul_ onwards, Eq. (22) can be analytically solved as well. We include full details on the derivation of this solution in Appendix B, but for brevity here we shall simply quote the result, which is

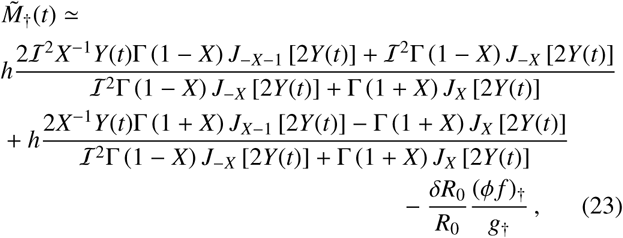

where one must define the following initial condition

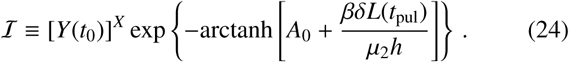

Using Eq. (23) in Fig. 4 we once again plot the approximative solution to the transmission dynamics against the full solution to the dynamics obtained numerically, each represented by the coloured solid and dashed lines, respectively, for a range of initial conditions *M* (*t*_0_) and two choices for *R*_0_ (separated by the left and right panels). The upper panels display the dynamics under the influence of a positive-valued migration-driven perturbation, and conversely, the lower panels display the dynamics under the influence of a negative-valued migration-driven perturbation.

**Figure 4.**
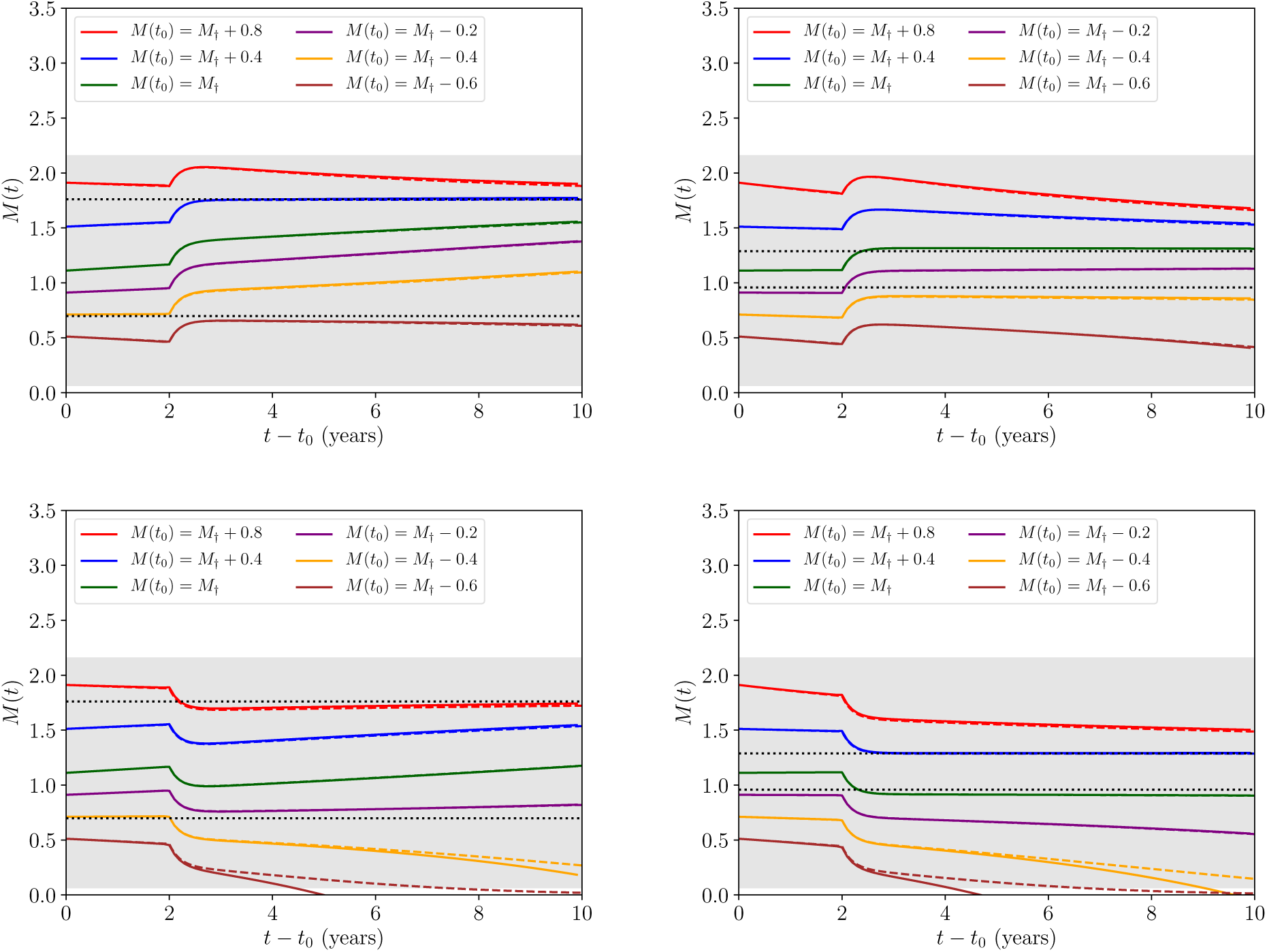
The approximative solution to the transmission dynamics given in Eq. (23) (applied where we have set *t*_pul_ = 2 years and Eq. (18) as the solution to the dynamics before this point) against the full solution to the dynamics obtained numerically, each represented by the coloured solid and dashed lines, respectively, for a range of initial conditions *M*(*t*_0_). All lines and the grey region correspond to their equivalent values in Fig. 2, where an additional upper and lower panel split in this set of figures compares the relative dynamical behaviour under the influence of a migration perturbation of *βδL*(*t*_pul_) = +1(−1) years^-1^for the upper (lower) pair of panels.

As in the case without migration, Fig. 4 demonstrates that the approximations made to obtain Eq. (23) only depart from the fully numerical solution in a significant way when the condition of expansion |*M* – *M* _†_(*z, k*)| < 1 (satisfied within the grey region) is no longer met. Comparing Fig. 2 with Fig. 4 also demonstrates very clearly how the long-time transmission dynamics of the system near the breakpoint are apparently quite unstable to an 𝒪(1) perturbation due to migration — even in the proximity of the ‘stable’ attractor equilibrium (upper of the two dashed horizontal black lines).

It is important to note here how, at much later times *t≫t*_0_, Eq. (23) is identical to Eq. (18) up to the inclusion of a modified initial condition *M*(*t*_0_) →*M*(*t*_0_)+*βδL*(*t*_pul_)/*µ*_2_. The simple form of this transformation, which arises from the large hierarchy in timescales, will motivate our Markovian model of stochastic migration in Sec. 4.

With our analysis in this subsection, we have begun to address the second of our two questions posed in the introduction. The main limitation of this approach so far is that we have only considered a single discrete migratory event. Therefore, in order to further evaluate the degree to which the unstable break-point and stable endemic equilibria are robust to the effects of migration and hence draw more concrete conclusions, we will need to develop a model of multiple migratory events. Before doing so we will need to introduce another expansion.

### 3.4. Expansions about the equilibria

In parallel with the expansion we have used previously, one may also derive an expansion around either the stable endemic equilibrium or unstable breakpoint equilibrium solutions *M* = *M*_*_(*z, k, R*_0_) to Eq. (9). This expansion takes the same form for both equilibria and is given by the following general expression

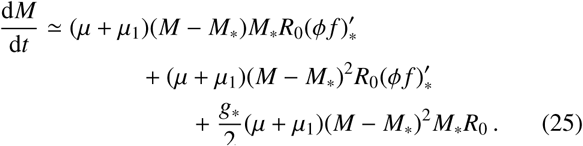

From Eq. (25) we hence obtain solutions analogue to Eq. (23), with the form

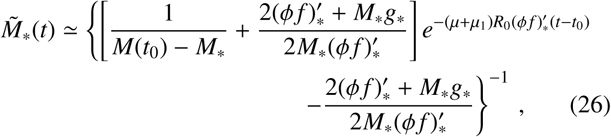

where we are making use of the familiar notation from earlier, (*ϕ f*) ≡ *ϕ*[*M*(*t*); *z, k*] *f* [*M*(*t*); *z, k*], such that

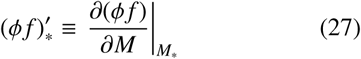

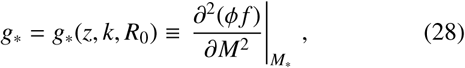

where we have now defined 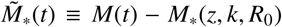. Depending on the choice of *M*_*_ (which corresponds to two degenerate values ∀*R*_0_ > *R*_0†_ as discussed previously), we may thus describe the transmission dynamics parameterically near to the stable or unstable equilibrium points, given a value for *R*_0_.

In Fig. 5 we plot this approximate solution given by Eq. (26) against the corresponding numerically-obtained solutions to Eq. (6) for a value of *R*_0_ = *R*_0†_(*z, k*) + 1.0 ≃ 3.08. Notice, in particular, how the timescale for the approximative expansion to break down is much shorter in the case of the unstable equilibrium — as we have only been able to plot up to 2 years before the diverging solutions completely leave the neighbourhood of *M*_*_ — which is due to the fact that the rate of time evolution in the transimission dynamics increases with increasing *R*_0_. This latter point becomes consistent with our expectations when noting that the amplitude of *M*″(*t*) as given by Eq. (8) grows (with a negative sign) with larger *R*_0_ values.

**Figure 5.**
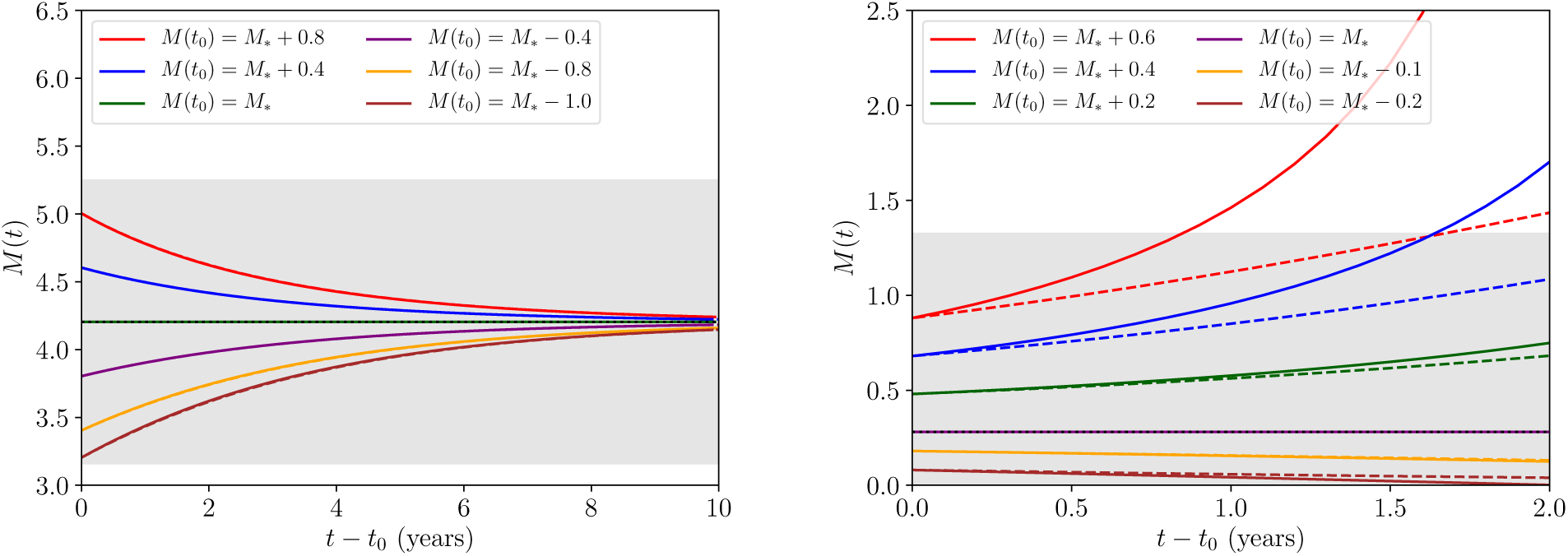
The approximative solution to the transmission dynamics given in Eq. (26) against the full solution to the dynamics obtained numerically, each represented by the coloured solid and dashed lines, respectively, for a range of initial conditions *M*(*t*_0_) and having fixed a value of *R*_0_ = *R*_0†_(*z, k*) + 1.0 ≃ 3.08. In the left panel we plot the transmission dynamics around the stable equilibrium, and in the right panel we plot the the dynamics around the unstable equilibrium. The dotted black horizontal line corresponds to the value of *M* = *M* *(*z,k,R*_0_) at the stable (left panel) and unstable (right panel) equilibrium numerically obtained from Eq. (9). Lastly, the grey region corresponds to values for which |*M* − *M* *(*z,k,R*_0_)| < 1.

Building from the formalism previously discussed, we may derive a theory of deterministic pulses around both equilibria as well, however the conclusions drawn from such an analysis would be the same as those of the previous section. Instead of such a theory, which has proven useful to analyse the stability of the system parametrically near the saddle-node bifurcation point, let us point out that in a slightly more realistic scenario: many such migratory events which perturb the reservoir may occur over the STH transmission period of interest. In the proceeding section we shall develop a theory which includes this possibility by utilising a stochastic process.

## 4. A stochastic theory of migration perturbations

### 4.1. Deriving the stochastic term

In this subsection we shall focus on obtaining a realistic stochastic representation of the flux in egg and/or larvae count into (or out of) the infectious reservoir over time. In order to do this, it shall serve our purpose to first briefly review the current theory regarding the probabilistic modelling for STH infections [6] and subsequently extend the approach to develop our model.

The well-known probabilistic derivation of the reservoir-driven *ϕ*(*M*; *z*; *k*) *f*(*M*; *z, k*) term, included in the force of infection of Eq. (1), involves indentifying the egg count e(*n*) generated by an individual as a function of the total number of worms *n* carried by that individual which includes both the density (aggregation of worms) dependence of fecundity (encoded by the *z* ≡ *e*^−*γ*^ parameter) and marginalising over the binomial probability of *n*_f_ of the worms being female,^7^ hence

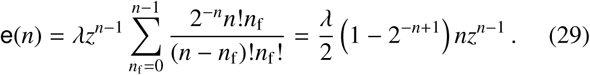

Taking the first moment of e → E(e) with respect to the known negative binomial probability mass function [6] of the total worm burden distribution within hosts

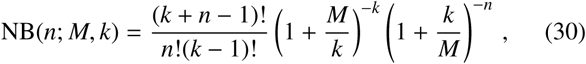

yields E(e) = *λϕ*(*M*; *z*; *k*) *f* (*M*; *z, k*)/2 ≡ *λ*(*ϕ f*)/2 by construction, where we shall continue to use this shorthand for brevity. The variance of e(*n*) may also be calculated as follows

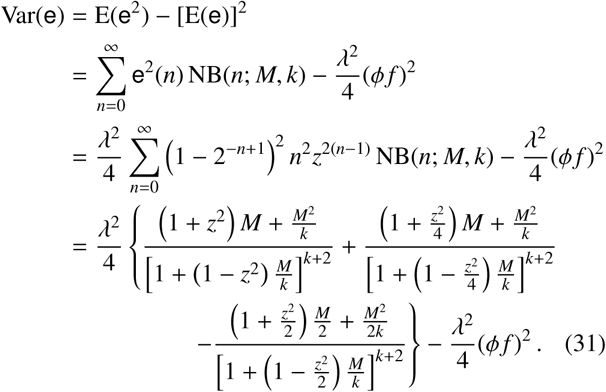

Although the moments of the egg count distribution are calculable, it is immediately unclear as to whether the shape of the egg count distribution inherits a negative binomial shape from the worm burden. By small argument expansion of Eq. (29) we find that in the limit where the total number of worms *n* ≪1/*γ* one can obtain the approximate relation

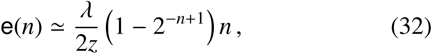

where, upon further restricting the number of worms to *n* ≳ 5, the 1 − 2^−*n*+1^factor in the expression above may be neglected to good approximation. This argumentation suggests that for numbers of worms in the range 1/*γ* ≫ *n* ≳ 5, or equivalently numbers of eggs in the range *λ*/(2*γz*) ≫ e ≳ 5*λ*/(2*z*), the egg count is effectively directly proportional to the total number of worms, and hence the distribution over the egg count may be well approximated by the following negative binomial shape

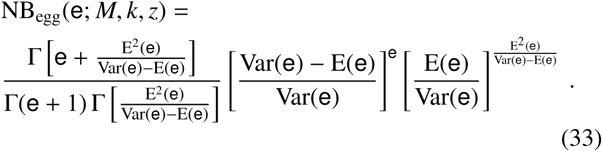

The event-based approach to a stochastic model of migration to or from the region of interest suggests that the times at which individuals do so are drawn from a Poisson distribution, and hence the event intervals are exponentially distributed

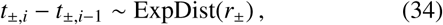

where *r*_±_ = *r*_+_ or *r*_−_, denoting the rates of people entering or leaving the region of interest, respectively — or, as we shall hereafter refer to them, ‘migratory rates’. Our proposed stochastic model is thus a compound Poisson process, with jump sizes e_*i*_ drawn from the egg count distribution that we argued for earlier in Eq. (33) and independently chosen, potentially time-dependent, parameters M = M(*t*) and k = k(*t*), such that

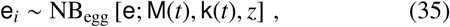

with a further normalisation included to account for the number of people within the region *N*_p_, such that our process *ℓ*_±_(*t*), which fluctuates around the deterministic dynamics in continuous time, becomes the following

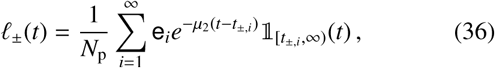

where 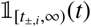 is an indicator function which is defined to take value 1 for arguments *t* ∈ [*t*_±,*i*_, ∞) and value 0 otherwise.^8^

In Eq. (36) the exponential death rate, which has been intuited from the functional form of Eq. (22), is a complicating factor that apparently renders this process non-Markovian for events separated in time by less than 1/*µ*_2_. Motivated by the simple form of pulse transformation (*M*(*t*_0_) → *M*(*t*_0_) + *βδL*(*t*_pul_)/*µ*_2_) in Sec. (3.3), we shall also consider a Markovian approximation to Eq. (36) which is made by temporally coarsegraining (integration over time) such that one obtains the following compound Poisson process in continuous time

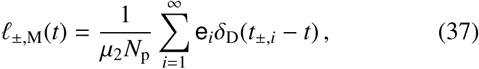

with an expected jump size and variance given by

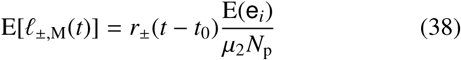

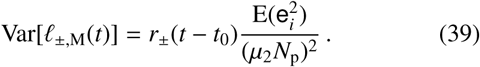

### 4.2. Computing the variability in the mean worm burden

Using Eq. (36), and taking note of the functional form given by Eq. (11), we write the following stochastic differential equation which combines both mean value drift and egg count distribution jumps which arise from the movements of people in and out of the given region of interest

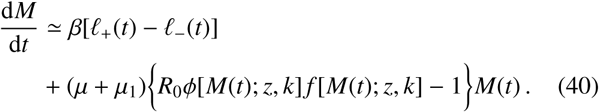

By computing numerical integrations over an ensemble of realisations for Eq. (40) and subsequently binning the samples at different ‘snapshots’ in time, one is able to construct a numerical approximation to the time evolution of the probability density function over the ensemble of possible worm burdens *P*(*M, t*).

In Fig. 6 we have initialised the *P*(*M, t*) distribution at the stable endemic (right column) and unstable breakpoint (left column) equilbrium values *P*(*M, t*_0_) = *δ*_D_(*M*_*_−*M*), where *M*_*_ here is the solution for either of the equilibria from Eq. (9), where we have set both *R*_0_ = *R*_0†_(*z, k*) + 0.1 ≃ 2.18 and the number of people *N*_p_ = 100. The solid lines indicate that the fully non-Markovian process given by Eq. (36) is used, whereas the dashed lines indicate the corresponding choice of approximate Markovian process, given by Eq. (37), which appears to match the fully non-Markovian process well.

**Figure 6.**
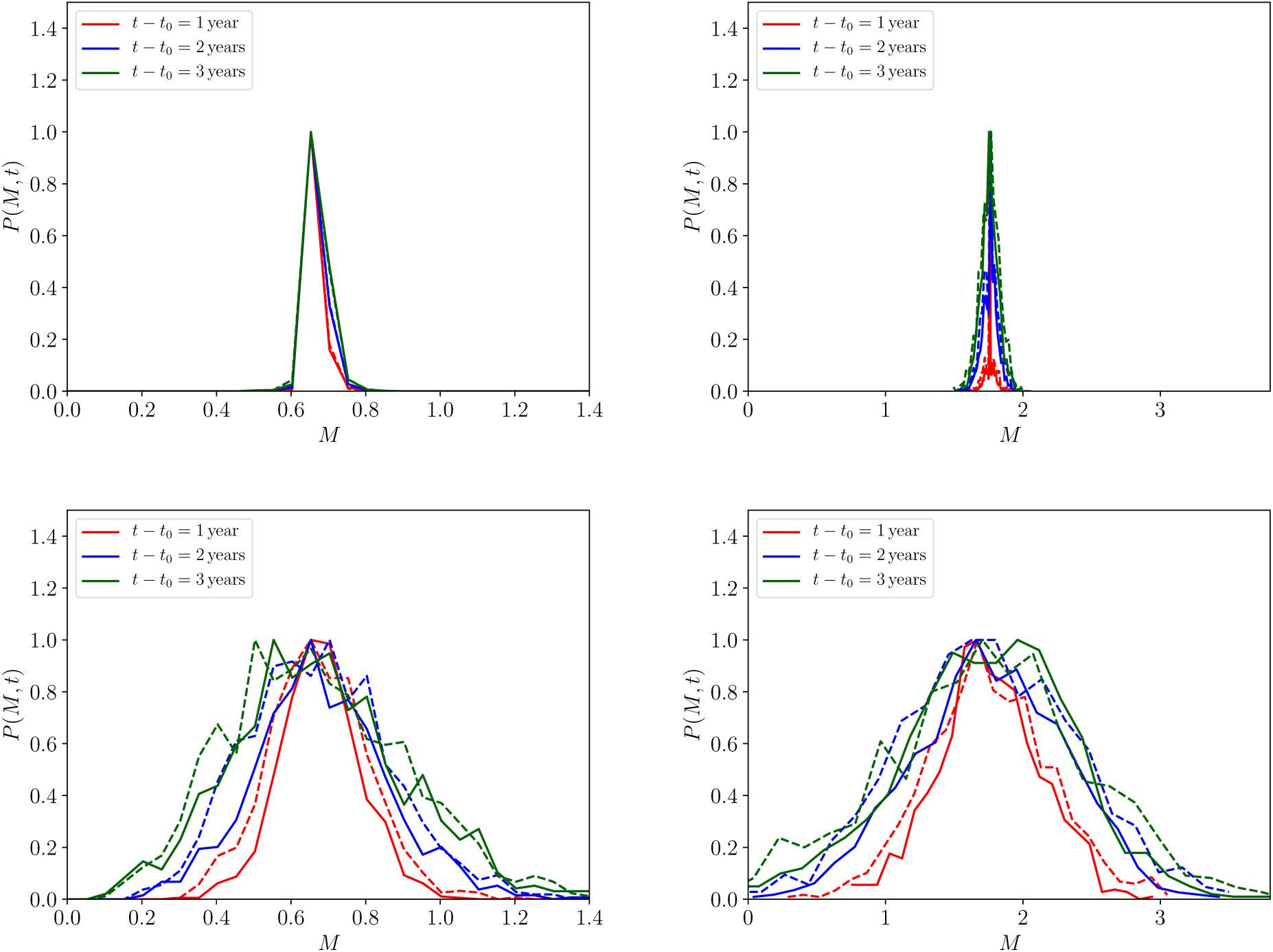
A comparison between the probability distributions *P*(*M, t*) at different snaphots in time (denoted by different colours) using *r* = 0.1*µ*_2_ (top panels) and *r* = 10*µ*_2_ (bottom panels) migration rates, having initialised *P*(*M, t*_0_) = *δ*_D_(*M*_*_ − *M*) (where *δ*_D_ is a Dirac delta function) at the unstable (left column) and stable (right column) equilibrium points *M*_*_, given a choice of *R*_0_ = *R*_0†_ (*z, k*) + 0.1 ≃ 2.18. The solid lines indicate that the fully non-Markovian process given by Eq. (36) is used, whereas the dashed lines indicate the corresponding choice of approximate Markovian process, given by Eq. (37). The distributions themselves have been numerically obtained by binning 10^3^realisations of Eq. (40).

For illustration purposes, in Fig. 6 we have also made the symmetric choice *r* = *r*_+_ = *r*_−_ as well as setting egg count distributions, which are given by Eq. (35), both with the same constant parameters, such that M(*t*) = *M*_*_ and k(*t*) = *k*. In the upper row of plots, we have plotted the distribution at different moments in time for a value of *r* = 0.1*µ*_2_ and, in the lower row of plots, a value of *r* = 10*µ*_2_, where it is immediately clear that a substantial variance is induced by migratory rates of *r* ≳ *µ*_2_.

Maintaining choices as above for Fig. 6, the full formula for the variance Var[*M*(*t*)] = E[*M*^2^*t*)] − {E[*M*(*t*)]}^2^, which is derived in detail in Appendix C in the limit of a Markovian process *ℓ*_±,M_(*t*), simplifies substantially to

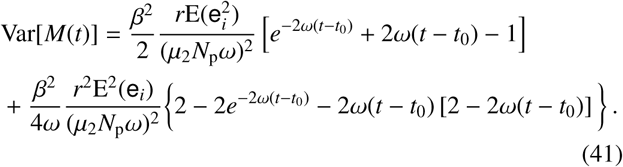

In Fig. 7 we plot the variances for the same system as before but with a wider range of values of the migratory rates *r*. We have provided the time evolution curves for the variance computed numerically for the fully non-Markovian process (solid coloured lines) and numerically for the Markovian process (dotted coloured lines), where the latter process is also computed with the analytic approximation provided by Eq. (41) (dashed coloured lines). The overall agreement between the numerically-obtained Markovian and non-Markovian processes is excellent — confirming the apparent agreement between the two distributions in Fig. 6 and strongly indicating that, due to the large hierarchy in dynamical timescales between the infectious reservoir and worm burden, migratory effects on the transmission dynamics appear effectively memoryless.

**Figure 7.**
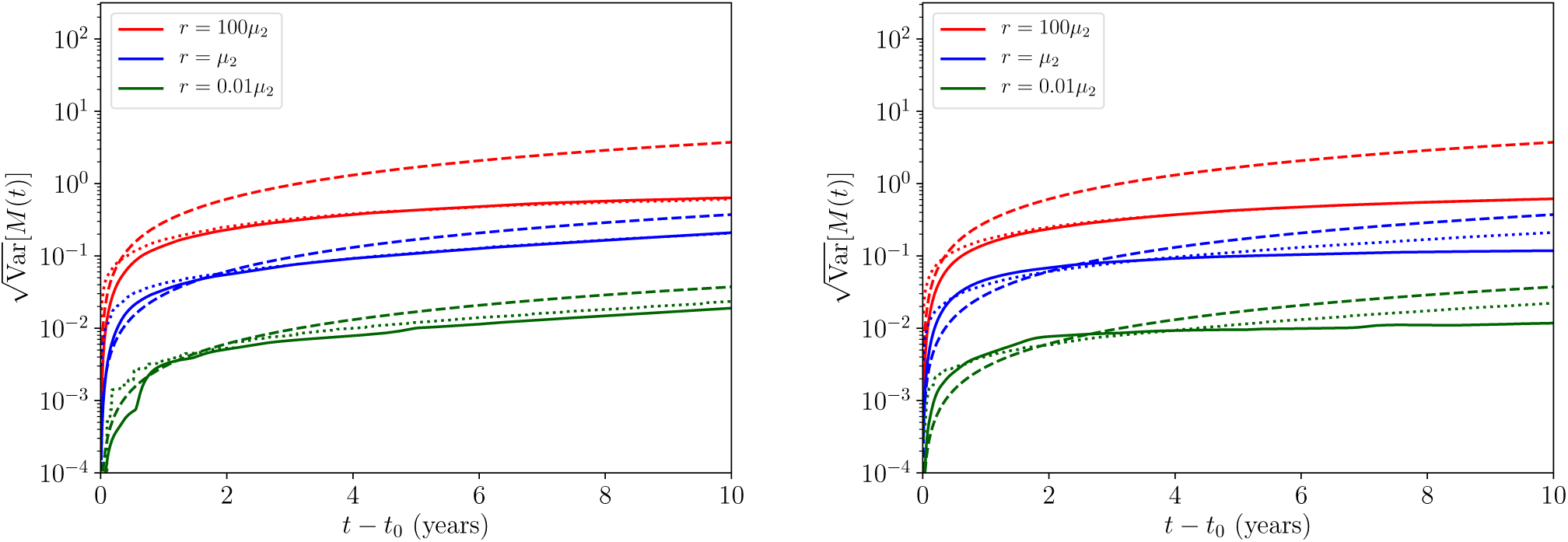
The root-variance over *M*(*t*) initialised at the unstable equilibrium *M*_*_ value (left panel) and at the stable equilibrium *M*_*_ value (right panel), plotted as a function of time, and numerically obtained by summing over 10^3^ realisations of Eq. (40) using the non-Markovian (solid coloured lines) and Markovian (dotted coloured lines) migration processes given by Eqs. (36) and (37), respectively. The dashed coloured lines correspond to the analytic Markovian solution given in Eq. (41).

The agreement between our analytic relation given by Eq. (41) and the numerical variances in Fig. 7 also appears to be good for a choice of the migratory rate *r ≲ µ*_2_. In the range where *r* > *µ*_2_, due to the disagreement between the numerical solution to the Markovian process and our approximation, we deduce that the total variance is limited by the presence of the disease extinction equilibrium *M*(*t*) = 0 — an effect which can already be observed for the tails of the distributions in Fig. 6.

We turn, finally, to readdressing the second of the two questions posed in the introduction. Our stochastic model of migratory perturbations has demonstrated that both the stable endemic and unstable breakpoint equilibria behave in a similar fashion under the influence of a symmetric migrating flux of people in and out of the infectious reservoir: when such a rate *r* is low when compared to the reservoir death rate *r ≪µ*_2_ the deterministic transmission dynamics experience some degree of controlled uncertainty; when *r≫ µ*_2_ our results indicate enormous variability in the dynamics may be potentially induced — as is indicated by the large variances illustrated in our Figs. 6 and 7.

## 5. Discussion

In this paper we have developed an analytic framework for describing the transmission dynamics of STH parasitic infections near the unstable equilibrium, or ‘breakpoint’ — the target for transmission elimination strategies using MDA and/or sanitation based programmes of soil-transmitted helminth control. Our analysis, in addition to extending the literature with new analytic insights, has also analytically and numerically investigated the effect that infected human migration can have on the dynamics of parasite transmission by developing the new models of discrete infectious reservoir ‘pulses’ and a stochastic theory of many migratory events. The tools we have developed, along with our specific findings, have important ramifications for both policies for control and the design of monitoring and evaluation programmes. As control efforts for neglected tropical helminth infections increase, our results should be applicable to many programmes today and in the future.

We assessed the rate of change in time for the transmission trajectories near the parasite transmission breakpoint (arising from the dieocious nature of helminth parasites), finding them to be extremely slowly varying. Illustrating this point quite clearly are our Figs. 2 and 3, where the timescales for significant change (and hence consequently of great relevance to monitoring and evaluating the impact of control programmes) can be of the order of many years. The slowly-varying dynamics around the unstable equilibrium thus represent a significant challenge to the detection of whether or not the breakpoint in transmission has been crossed and the STH population is on the way to extinction. Stochastic noise arising from low counts of infected people will further accentuate this detection challenge, since random variation may either lead to quick elimination or very drawn out trajectories to elimination.

A secondary, yet no less important effect, was studied in conjunction with our earlier analysis and is perhaps best illustrated by the combination of Figs. 6 and 7. The migration of infected humans into an area where control measures have driven STH prevalence below the transmission breakpoint was found to introduce significant uncertainty in the transmission dynamics near both the stable endemic and the unstable ‘breakpoint’ equilibrium.

In the limit where the migratory rate — the rate of people leaving and entering the region or community and introducing material into the infectious reservoir (eggs or larvae, depending on the species of STH) per unit time — is much smaller than the rate of reservoir death (the rate at which the eggs or larvae die outside of a host), our results indicate that uncertainty in the dynamics is present but controlled to some degree such that elimination will eventually occur. In the opposite limit, where the average migratory rate greatly exceeds the average death rate of infectious eggs or larvae in the reservoir, our results suggest that great variability in the transmission dynamics is possible and, indeed, likely. We therefore find that infected human migration is an important effect to quantify in order to assess its ability to undermine targets for parasite elimination from control measures, such as MDA and/or improvements in water and sanitation provision.

In addition to these important practical implications which may stem from human migration, our analysis has also yielded further insights into the theory of human helminth parasite transmission dynamics. The large differences in dynamical timescales present between the infectious reservoir or vector hosts in the case of schistosomes, filarial worms (life expectancy a few weeks to months) and adult worms in the human host (one to many years) admits an accurate Markovian description to be formulated for the migration model. This in turn provides analytical insights into the key parameters that determine epidemiological outcomes within the dynamic transmission systems.

Throughout this paper we have deliberately excluded the possible complicating effects induced by the presence of an age-structured population. Many components of our analysis should be generalisable in this aspect and we leave this interesting extension to future work.

## Acknowledgements

RJH, JET and RMA all gratefully thank the Bill and Melinda Gates Foundation for research grant support via the DeWorm3 (OPP1129535) award to the Natural History Museum in London (http://www.gatesfoundation.org/). CV gratefully acknowledges funding from the NTD Modelling Consortium (OPP1184344) by the Bill and Melinda Gates Foundation (http://www.gatesfoundation.org/) in partnership with the Task Force for Global Health (http://www.taskforce.org/). The views, opinions, assumptions or any other information set out in this article are solely those of the authors. All authors acknowledge joint Centre funding from the UK Medical Research Council and Department for International Development.

### Appendix A. The saddle-node bifurcation expansion in more detail

One can ‘prove’ that there is indeed a saddle-node bifurcation at the critical value where the stable and unstable equilibria collide. Taking Eq. (6), the Jacobian of the *M′* (*t*) = *F*[*M*(*t*)] one-dimensional system (i.e., ℐ = *F′* (*M*)) is

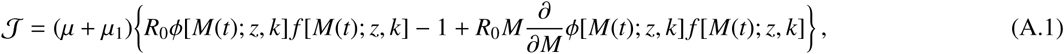

Along either of the equilibria *M*\*, Eq. (9) is satisfied, and so according to Eq. (A.1) one finds a singularity (ℐ = 0) when

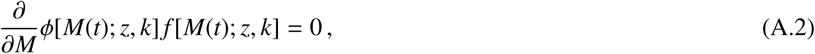

at which point a saddle-node bifurcation is known to occur. This point marks the collision between the stable endemic equilibrium and unstable equilibria and is known in the epidemiological literature as the disease ‘breakpoint’. ^9^ The value of *M* = *M*_†_(*z, k*) which satisfies the above relation can also be deduced

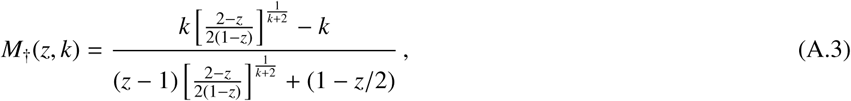

and the critical basic reproduction number *R*_0†_, which we introduced in Sec. 1, is thus clearly

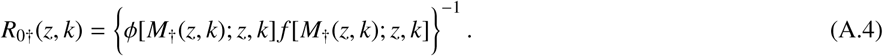

Given the existence of this saddle-node bifurcation, locally about *M*_†_(*z, k*) we may expand the system (suppressing dependencies and defining (ϕ *f)*_†_ ≡ ϕ[*M*_†_(*z, k*); *z, k*] *f* [*M*_†_(*z, k*); *z, k*] for brevity) such that

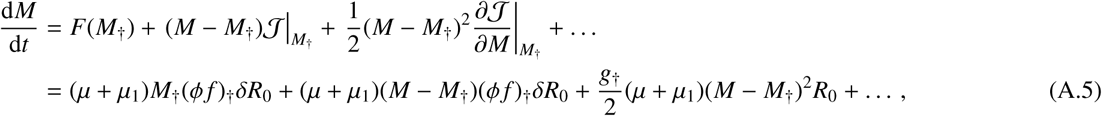

where we have defined *δ R*_0_ = *δR*_0_(*z, k, R*_0_) ≡ *R* _0_ – *R*_0†_(*z, k*) and a new function

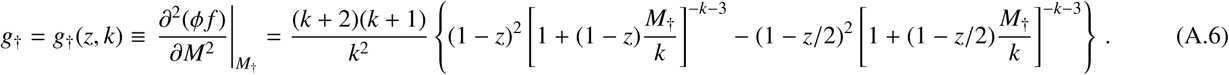

Note that, for typical values of *z*, the *g*_†_(*z, k*) function takes negative values which decrease in magnitude with increasing *k*.

Now by truncating the expansion in Eq. (16), keeping up to (𝒪*M*^2^) terms,^10^ we are able to solve the system exactly to find the following analytic solution

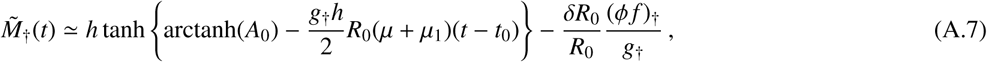

where 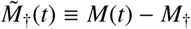 and we have defined

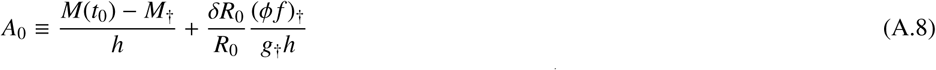

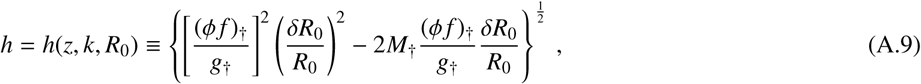

and one predictably finds that the latter function becomes imaginary for a finite *k* exactly when *R*_0_ becomes less than the critical value, i.e., *R*_0_ < *R*_0†_(*z, k*). As a consistency check, one may further confirm that this switch maps the hyperbolic tangent function Eq. (18) onto a standard tangent — a switch between hyperbolic to elliptic local geometries — and, consequently, the dynamics of *M*(*t*) switch generally from a possible endemic equilibrium attraction to pure decay *M*(*t*) → 0, ∀*M*(*t*_0_), as expected.

### Appendix B. Solving the system with a migration perturbation

In the main text, by truncating an expansion of Eq. (11) and keeping up to 𝒪(*M*^2^) terms in same way as in Eq. (16), we find that the resulting approximate equation takes a Riccati form

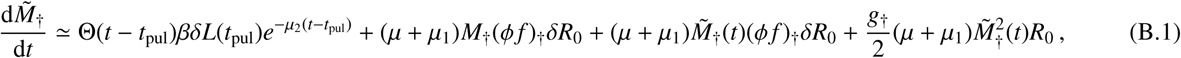

where, once again, 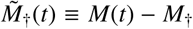 and Θ(*t – t*_pul_) is the Heaviside (step) function which initialises the migration pulse at time *t*_pul_ and *δL*(*t*_pul_) is the maximal amplitude of the perturbation in the infectious reservoir.

Remarkably, evolving the elapsed time from *t*_pul_ onwards, Eq. (22) can be analytically solved as well. If one picks the transformation

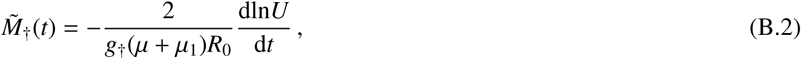

we may rewrite Eq. (22) as the following Sturm-Liouville equation

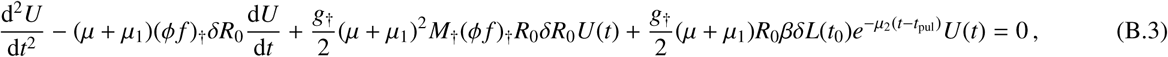

whose formal solution is known to be

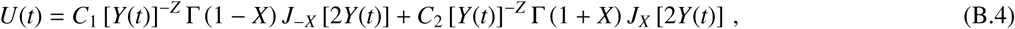

where *J*_*x*_(*y*) is the Bessel function of the first kind and, for brevity, we have have also defined

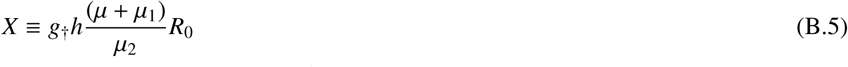

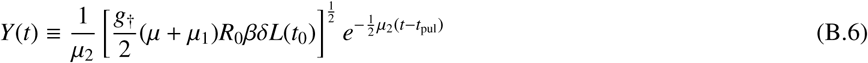

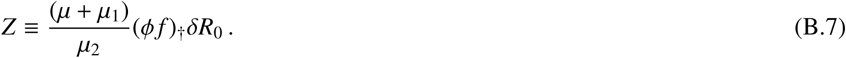

Using Eq. (B.2) to transform Eq. (B.4) back into into 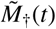 (and hence *M*(*t*)), we arrive at our general system solution after some straightforward manipulations

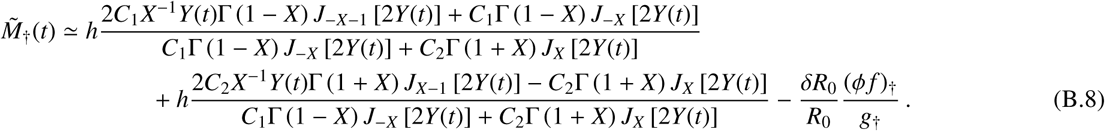

Notice that in the limit *δL*(*t*_0_) → 0 the solution above should asymptotically match the time dependence of Eq. (18). Taking this limit for Eq. (B.8) yields the following asymptotic behaviour

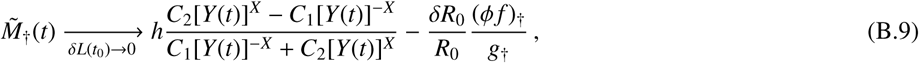

hence, by assuming that the initial conditions of Eq. (B.8) have no migration perturbation already, we match the two solutions with following choices for the constants^11^

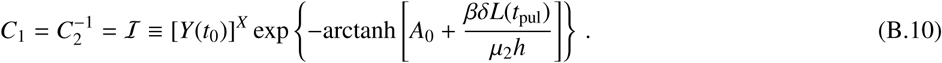

Therefore, the fully-determined solution corresponding to the approximative transmission dynamics in this case is reduced to

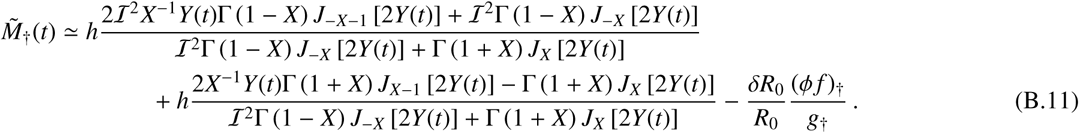

### Appendix C. Computing the variance over the ensemble of possible worm burdens

Motivated by the expansions in previous sections, we may take a linear approximation of Eq. (40) in the neighbourhood of one of the equilibria *M*_*_ in order to write the following approximate equation

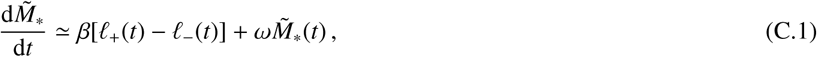

where 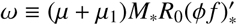 and we are reminded that 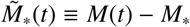. Eq. (C.1) admits the following implicit solution

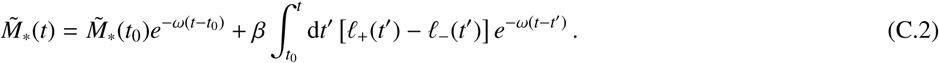

Let us now assume to be in the limit where *r* ≪ 1/*µ*_2_, hence we may replace the noise term with an approximately Markovian one ℓ_±_(*t*) → ℓ_±,M_(*t*) as in Eq. (37). Taking the ensemble average over separate realisations of Eq. (C.2), and initialising all realisations at some value *M*(*t*_0_), we thus arrive at the following solution for the first moment of *M*(*t*)

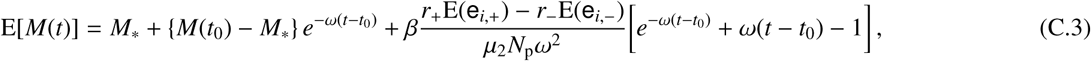

where we have made use of Eq. (38) and have included the possibility that two different egg count distributions — with the form of Eq. (35) — may be used to draw the jump sizes from for the ingoing e_*i*,+_ and outgoing e_*i*,-_ migrant patterns.

By using the Markov condition for the ℓ_±,M_(*t*) processes (where δ_D_ is a Dirac delta function)

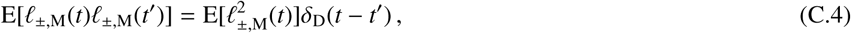

as well as assuming that no cross-correlations exist, such that E[ℓ_±,M_(*t*) ℓ_∓,M_(*t*′)] = 0, we may further use Eq. (C.2) to obtain the second moment of *M*(*t*), which is

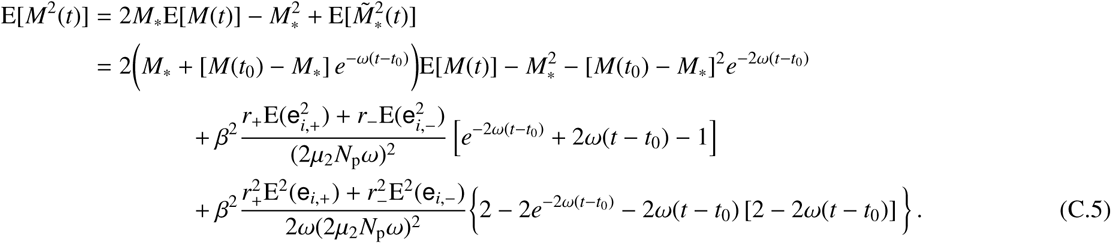

## Footnotes

1 Note that this is not the ‘true’ *R*_0_, which itself is not calculable from the next-generation method — this is because the spectral radius of the Jacobian for the force of infection in Eq. (2) vanishes for *M* = 0.

2 The Brent method is necessary in order to solve for the unstable equilibrium in particular, due to the convexity of the function one must minimise in this parameter regime.

3 This assumption is consistent with the timescale of reservoir infectious material death, *µ*_2_, being much shorter than the other two decay timescales *µ* and *µ*_1_.

4 Or, equivalently, in the neighbourhood of the saddle-node bifurcation the system becomes topologically equivalent to *M*′(*t*) = *a* + *bM*(*t*) + *cM*^2^(*t*).

5 The validity of this condition may be examined by finding the value of |*M*−*M*_†_(*z, k*)| at which the higher-order terms in our expansion in Eq. (16) become equal to the second-order term. We examine the ramifications of this condition in more detail in a future article [16].

6 The method used is a Runge-Kutta (4)5 method developed by Dormand & Prince, 1980 [17], and described in Ref. [18], where the `scipy.integrate`module in `python` provides a simple software implementation.

7 Note that, as in the standard result, this derivation assumes the total polygamy of male worms. Furthermore, the *n* − 1 summation limit arises from lack of generated eggs if the totality of worms within a given host are female.

8 This is equivalent to the Heaviside function earlier.

9 In more detail, this breakpoint arises due to a combination of helminth sexual reproduction and their the mating habits which allow for the existence of a threshold — explicitly here *M*_†_(*z, k*) — below which the disease may become dynamically driven to exinction for a finite initial condition in *M*.

10 Or, equivalently, in the neighbourhood of the saddle-node bifurcation the system becomes topologically equivalent to *M′*(*t*) = *a* + *bM*(*t*) + *cM*^2^(*t*).

11 There is a subtlety here in the choice of initial condition: due to the discontinuity at the boundary, an additio<u>na</u>l component of 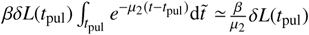 must be added to the initial worm burden from the infectious reservoir pulse, such that 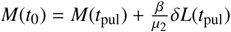.

